# FcγRIIB I232T polymorphic change allosterically suppresses ligand binding

**DOI:** 10.1101/586487

**Authors:** Wei Hu, Yong Zhang, Xiaolin Sun, Liling Xu, Hengyi Xie, Zhanguo Li, Wanli Liu, Jizhong Lou, Wei Chen

## Abstract

FcγRIIB bindings to its ligand suppress immune cell activation. A single-nucleotide polymorphic (SNP) change, I232T, in the transmembrane (TM) domain of FcγRIIB loses its suppression function, which clinically associates with systemic lupus erythematosus (SLE). Previously, we reported that I232T tilts FcγRIIB’s TM domain. In this study, combining with molecular dynamics simulations and single-cell FRET assay, we further revealed that such tilting by I232T unexpectedly bends the FcγRIIB’s ectodomain towards plasma membrane to allosterically impede FcγRIIB’s ligand association. We then used single-cell biomechanical assay to further find out that I232T also reduces two-dimensional in-situ binding affinities and association rates of FcγRIIB interacting with its ligands by three-folds. This allosteric regulation by a SNP provides an intrinsic molecular mechanism for functional loss of FcγRIIB-I232T in SLE patients.

## Introduction

Disorders or hyper activation of immune components could lead to autoimmune diseases. Malfunction of an immune receptor, FcγRIIB, is generally regarded as destructive for immune system ^1-3^. FcγRIIB is widely expressed on most types of immune cells including B cells, plasma cells, monocytes, dendritic cells, macrophages, neutrophils, basophils, mast cells and even memory CD8 T cells ^4^. FcγRIIB is unique among all immune-receptors for Fc portion of IgG molecules (FcγRs), which efficiently down-regulates the activation of immune cells. It has been shown that single nucleotide polymorphisms (SNPs) of the human FcγRIIB gene extensively influence the susceptibility towards autoimmune disorders ^2-3,5^. A T-to-C variant in exon 5 (rs1050501) of FcγRIIB causes the I232T substitution (FcγRIIB-I232T) within the transmembrane (TM) domain, and positively associates with systemic lupus erythematosus (SLE) in the homozygous FcγRIIB-I232T populations through a large amount of epidemiological studies ^2,5-9^. Although a statistical linkage of the homozygous FcγRIIB-I232T polymorphism with SLE was established, comprehensive assessments and deeper mechanistic investigations towards the inter-linkage of FcγRIIB-I232T regarding to the age of syndrome onset, progress, and clinical manifestation of SLE are still lacking.

## Results and Discussion

In this report, we firstly performed systemic examination over the association of FcγRIIB-I232T with clinical manifestations of SLE. We enrolled 711 unrelated Chinese patients with SLE and complete clinical documents into this study (Table 1). 688 unrelated healthy Chinese volunteers with matched gender and age were then enrolled as controls. We confirmed a strong positive association of the homozygous FcγRIIB-I232T polymorphism with SLE (*χ*^*2*^ = 7.224, *p* = 0.008, odds ratio with 95% confidence interval (CI) = 1.927) (Table 1), in consistent with the published epidemiological data ^2^, ^5-9^. Next, we comprehensively analyzed the clinical data for all 711 SLE patients, including 50 FcγRIIB-I232T homozygotes, 283 FcγRIIB-I232T heterozygotes and 378 FcγRIIB-WT carriers. Strikingly, we found that the homozygous FcγRIIB-I232T polymorphism is significantly associated with early disease onset (age at disease onset < 37, *p* = 0.002) (Supplementary file 1 and 2). We also observed a significant association of the homozygous FcγRIIB-I232T polymorphism with more severe SLE clinical manifestations since the corresponding SLE patients present significant elevation in the amounts of anti-dsDNA antibodies (*p* = 0.004), anti-nuclear antibodies (*p* = 0.021) and total Immunoglobulin (Ig) (*p* = 0.032) when compared to patients carrying heterozygous FcγRIIB-I232T polymorphism or FcγRIIB-WT (Supplementary file 1 and 2). Moreover, homozygous FcγRIIB-I232T polymorphism is also significantly associated with the higher SLE disease activity index (SLEDAI) score (*p*= 0.014 for SLEDAI ⩾12 vs. *p* = 0.861 for SLEDAI < 12) as well as more severe clinical manifestations including arthritis (*p* = 0.008), anemia (*p* = 0.006), leukopenia (*p* = 0.005), complement decrease (*p* = 0.006), hematuria (*p* = 0.004) and leucocyturia (*p* = 0.010) (Supplementary file 1 and 2). A suggestive association was also observed between homozygous FcγRIIB-I232T polymorphism and serositis (*p* = 0.063) (Supplementary file 1 and 2). These association analyses demonstrated that SLE patients homozygous for FcγRIIB-I232T polymorphism are prone to develop more severe clinical manifestations than the patients carrying heterozygous FcγRIIB-I232T polymorphism or FcγRIIB-WT, reinforcing the importance to study the pathogenic mechanism of FcγRIIB-I232T polymorphism since this SNP occurs at a notable frequency in up to 40% (heterozygous polymorphism) humans ^2-3^.

**Table 1:** Association analysis of rs1050501 with SLE (adjusted for sex and age)

| rs1050501 | Control | SLE | OR | 95% CI | *p* value |
| --- | --- | --- | --- | --- | --- |
| **allelic** |  |  |  |  |  |
| T | 1038 (75.4) | 1039 (73.1) | 1.142 | 0.958-1.362 | 0.138 |
| C | 338 (24.6) | 383 (26.9) |  |  |  |
| **genotypic** |  |  |  |  |  |
| TT+TC | 662 (96.2) | 661 (92.7) | 1.927 | 1.185-3.134 | 0.008 |
| CC | 26 (3.78) | 50 (7.03) |  |  |  |

Previous biochemical studies revealed that monocytes harboring FcγRIIB-I232T are hyper-activated with augmented FcγRI-triggered phospholipase D activation and calcium signaling ^10^. B lymphocytes expressing FcγRIIB-I232T are of hyperactivity and abnormal elevation of PLCγ2 activation, proliferation and calcium mobilization ^11^. FcγRIIB-I232T B cells lose the ability to inhibit the oligomerization of B cell receptors (BCRs) upon co-ligation between BCR and FcγRIIB ^12^. Recent live-cell imaging studies showed that B cells expressing FcγRIIB-I232T fail to inhibit the spatial-temporal co-localization of BCR and CD19 within the B cell immunological synapses ^13^. Human primary B cells from SLE patients with homozygous FcγRIIB-I232T mutation revealed hyper-activation of PI3K ^13^. Thus, FcγRIIB-I232T is very likely the first example that a naturally occurring SNP within the TM domain of a single-pass transmembrane receptor can cripple its function in principle and is significantly relevant in diseases.

These signaling events are usually triggered or followed by ligand engagement of FcγRIIB, while this function is disrupted by a single amino acid change from Ile to Thr in the TM domain. Two early biochemical studies proposed a model of reduced affinity between FcγRIIB-I232T and lipid rafts to explain the functional relevance and effect of this mutation ^10-11^. Another model suggested that I232T mutation enforces the inclination of the TM domain and thereby reduces the lateral mobility and inhibitory functions of FcγRIIB. However, both models assumed that FcγRIIB-I232T and FcγRIIB-WT (I232) have an equal capability to perceive and bind to the ligand, the IgG Fc portion within the antibody antigen immune complexes. This important but experimentally un-proved pre-requisition in both models is based on the argument that FcγRIIB-I232T and FcγRIIB-WT (I232) are identical in terms of the amino acid sequences of the extracellular domain and thus the quaternary structures for recognizing the ligands, i.e., the IgG Fc portions ^14-15^. However, currently there are no experimental evidences to validate this pre-requisite assumption.

We thus investigated whether I232T polymorphic substitution in the TM domain of FcγRIIB allosterically affects ligand recognition. Our previous observation of the forced inclination of TM domain by I232T led us to hypothesize that the inclination of TM domain may lead to ectodomain conformational changes to allosterically attenuate ligand binding. We first carried out large-scale molecular dynamic simulations with full human FcγRIIB imbedded in the lipid bilayer harboring residues I232 or T232 on its TM domain (Figure 1A & Figure 1—figure supplement 1A). The simulations confirmed previous results with TM only^16^, *i.e.*, I232T polymorphic substitution enforces the inclination of the TM domain (Figure 1B, right). The inclination might be resulted by the ability of H-bond formation between the side-chain Oγ atom of T232 and the backbone oxygen atoms of the neighboring residues in T232 system (Figure 1B, left). The differences on the orientation of the TM domain induce a different conformation on the membrane proximal region (ecto-TM linker) at the extracellular side (Figure 1—figure supplement 2). The membrane buried non-helical region of the linker extends more in the I232T form than that in the WT, and the length between S218 and P221 peaks at 11 Å for I232T, 3 Å longer than the 8 Å peak position for I232 system (Figure 1C). This length elongation further results in a different conformation of residue P217, the main chain dihedral angle of P217 in I232 system displays two populations at 141°±23° and -50°±12°, respectively, but shifts to 4°±45° and -75°±12° in the T232 form (Figure 1D and Figure 1—figure supplement 2). These effects propagate and lead to striking effect on the extracellular domains of FcγRIIB. We found that the extracellular domain of the T232 form adopts significant different conformation than that of the I232 form. The ectodomain of I232 maintains more straight conformation, whereas that of T232 bends down towards the lipid bilayer (Figure 1E). Statistical analyses show that the ectodomain inclination angle of the T232 form distributes across 30∼60° with a sharper single-peak at 40° (Figure 1E). In contrast, the angle of the I232 form distributes more flatter with a most favorable probability ranging from 50° to 70° (Figure 1E). The distance of C1 domain is much closer to the membrane for the T232 form than the I232 form (Figure 1E). These results suggest that the T232 morphism (or I232T mutation) may reduce the antibody recognition ability of FcγRIIB via two aspects. First, although the Fc binding site is not buried, the orientation and membrane binding of T232 may sterically prevent the accessibility of the Fc portion of IgG, as significant clashes between docked Fc and the membrane are observed (Figure 1—figure supplement 1B). Second, T232 is more rigid (or less flexibility, Figure 1E) such that the chance to associate with the ligand is decreased (thus the ligand association rate may be significantly reduced).

**Figure 1.**
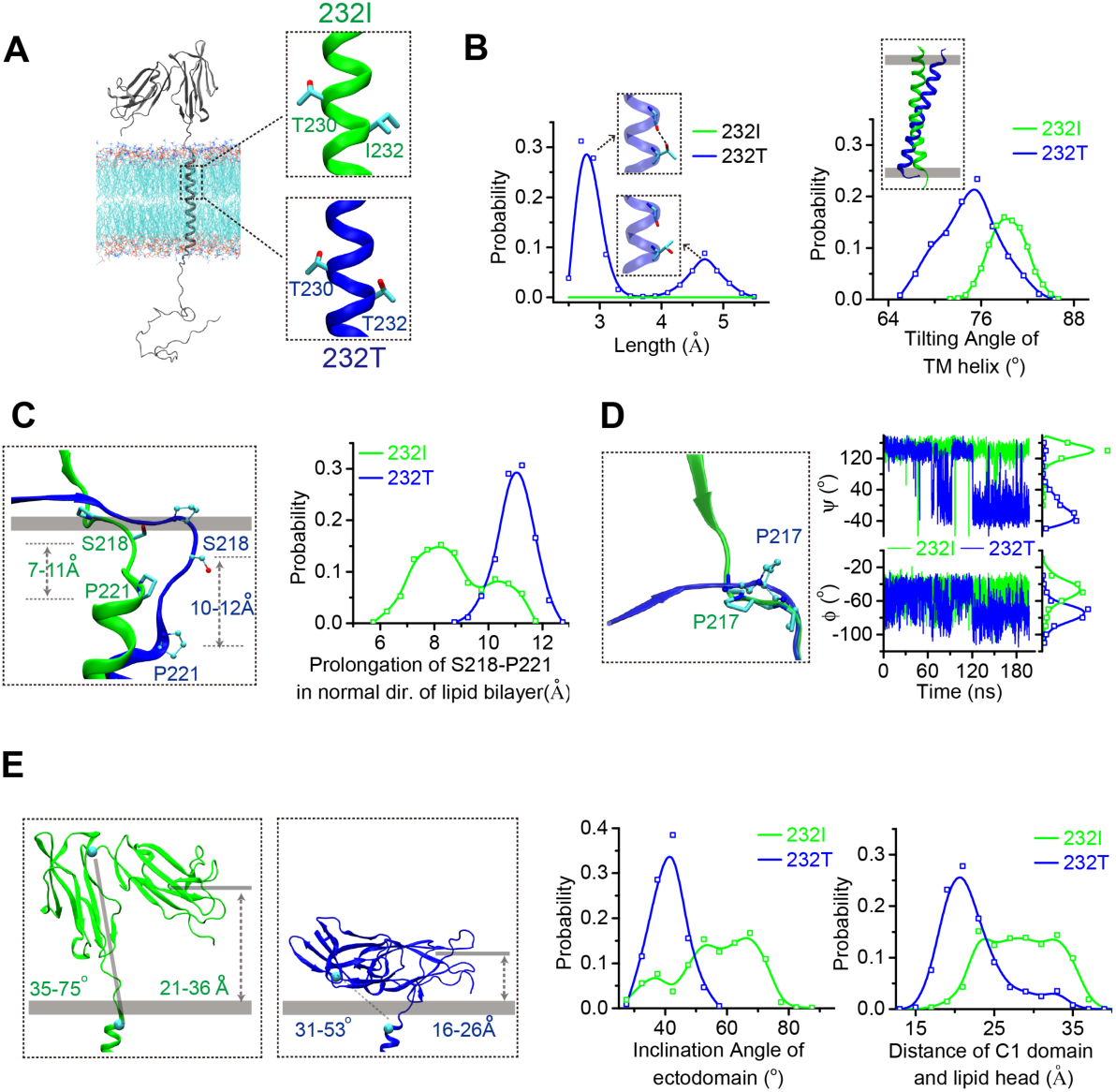
MD simulations reveal the conformational dynamics of WT FcγRIIB and its I232T polymorphic change. (A) The modeled structure of complete FcγRIIB (residues A46-I310, shown in grey cartoon) imbedded in an asymmetric lipid bilayer (Connected lines with atoms colored by type: P, tan; O, red; N, blue; C, cyan). The helical structures in the vicinity of residue 232 for WT (I232, green) and I232T (T232, blue) are shown in the insets. (B) Probability distributions of the distance between T232 Oγ atom and its nearest backbone O atom from residue V228 (left), and of the tilting angles between TM helix and lipid bilayer (right), the inclination of TM for T232 can be observed clearly. (C) Comparison of the representative snapshots of I232 and T232 systems at the stalk and TM linker region after superposing the lipid bilayers (left), and length distribution of S218-P221 backbone in normal direction of lipid bilayer (right). (D) Conformational difference of I212-S220 regions after aligning residues S218 to S220 (left), and the time courses of the dihedral angles ψ, φ) of residue P217 (right). (E) Representative snapshots of the I232 and T232 systems with the inclination angles and C1(Ig-like C2-Type 1 domain)/bilayer distances indicated. Probability distributions of the inclination angle between FcγRIIB ectodomain and lipid bilayer (left), and the distances between C1 domain and lipid bilayer (right).

MD simulations suggest that FcγRIIB ectodomain may bend towards membrane through weakly association of its ectodomain with the membrane via multiple sites (Figure 1—figure supplement 3) for I232T polymorphism. We next performed single-cell fluorescence resonance energy transfer (FRET) assay to experimentally validate whether I232T polymorphism allosterically bends the FcγRIIB ectodomain towards cell membrane (Figure 2A). According to our MD simulation results (Figure 1E), we hypothesized that an mTFP (as FRET donor) fused at the N-terminal of FcγRIIB (232I or 232T) ectodomain should fall in the spatial proximity (∼16∼36Å) for FRET with plasma outer membrane labeled with octadecyl rhodamine B (R18, as FRET acceptor), and that I232T polymorphism may enhance FRET efficiency. With de-quenching assay on A20II1.6 B cell lines expressing similar level of either mTFP-232I or mTFP-232T FcγRIIB (Figures 2B and 2C), we found that I232T polymorphism indeed enhances the FRET efficiency about two folds, from ∼ 20% in the I232 form to ∼40% in the T232 form (Figure 2C and 2D). This enhancement of FRET efficiency by I232T polymorphism indicates that FcγRIIB-232T ectodomain prefers to a more recumbent orientation on the plasma membrane than FcγRIIB-232I, consistent with our MD simulation observations above.

**Figure 2.**
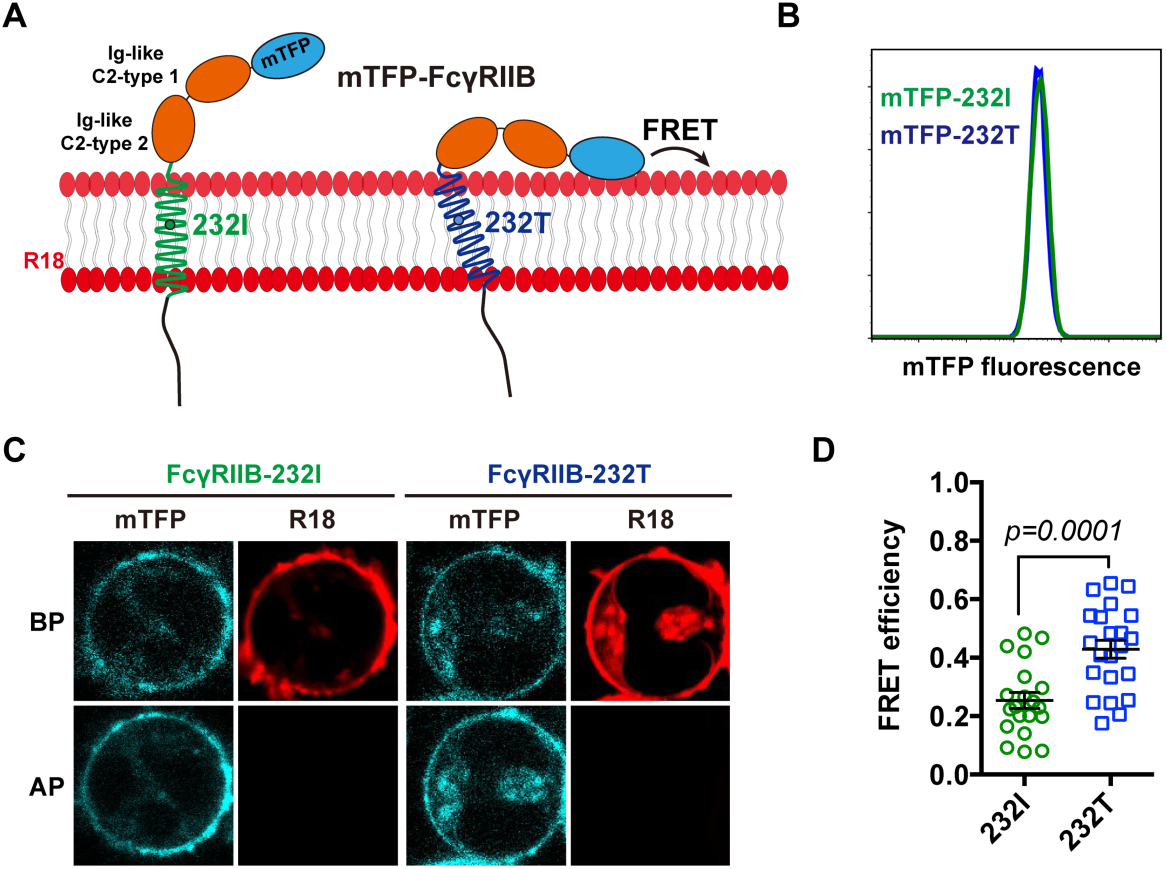
The FcγRIIB-232T ectodomain prefers to a more recumbent orientation on the plasma membrane. (A) Schematic illustration of mTFP-R18 FRET system to detect the distance between the ecto-domain of FcγRIIB232I (green) or FcγRIIB232T (blue) (N-terminal of ectodomain fused with mTFP as FRET donor, cyan) with the plasma membrane (stained with R18 dye as FRET acceptor, red). (B) mTFP fluorescence comparison of A20II1.6 B cell lines expressing mTFP-232I (green) or mTFP-232T (blue) constructs by FACS analysis. (C) Representative images of de-quenching FRET assay. R18-labeled mTFP-232I or mTFP-232T cells image were acquired in both channels before or after R18 photo-bleaching (BP or AP). (D) FRET efficiency of mTFP-232I or mTFP-232T cells (∼20 cells, respectively) were calculated and plotted. Error bars represent SEM.

Ectodomain orientation changes of a receptor can significantly affect its *in-situ* binding affinity with its ligands^17^. We therefore predict that titling FcγRIIB ectodomain towards plasma membrane by I232T polymorphism may attenuate its ligand binding affinity, especially the association rate. To test this hypothesis, we applied well-established single-cell biomechanical apparatus with adhesion frequency assay^18^ to directly and quantitatively measure *in-situ* two-dimensional (2D) binding kinetics of the WT or I232T FcγRIIB binding with its ligands (Figure 3A). It revealed that the *insitu* 2D effective binding affinity of FcγRIIB232I with MERS virus S protein human IgG1 antibody (anti-S) is about three times higher than that of FcγRIIB232T binding with same antibody (*A*_c_*K*_a_=3.03±0.15×10^-7^ and 0.80±0.04×10^-7^µm^4^, respectively), whereas that of human IgG4 is hardly measured as its binding is too weak and beyond the detection limit (10^-8^ µm^4^) ^18^ of this assay (Figure 3B and 3D), which is consistent with previous reported FcγRIIB/IgG4 binding affinity is far less than IgG1 ^1^. Moreover, off-rates of the WT and the I232T form binding with human IgG1 are similar (7.75±1.42 and 7.62±1.41 s^-1^, respectively) (Figure 3B and 3F), while the 2D effective on-rate of the I232T binding with human IgG1 is three times slower than that of the WT binding with same ligand (Figure 3B and 3E). These kinetics data strongly support our prediction that I232T polymorphism tilts FcγRIIB ectodomain more recumbent toward the plasma membrane so that its ligand binding domain is harder to be accessed, which reduces FcγRIIB/IgG1 binding on-rate. The conclusion is also confirmed by FcγRIIB binding with another human IgG1 (HIV1 gp120 human IgG1, anti-gp120) (Figures 3C to 3F). That is, the 2D effective affinity of FcγRIIB binding with human IgG1 and on-rate both are three times higher than those of 232T’s (*A*_c_*K*_a_=7.74±0.24×10^-7^ and 2.43±0.11×10^-7^µm^4^, respectively; *A*_c_*k*_on_=5.95±0.19 and 2.16±0.10×10^-7^µm^4^ s^-1^, respectively), while their binding off-rates are similar (7.70±0.83 and 8.90±1.61 s^-1^, respectively) (Figure 3C, 3D, 3E and 3F).

**Figure 3.**
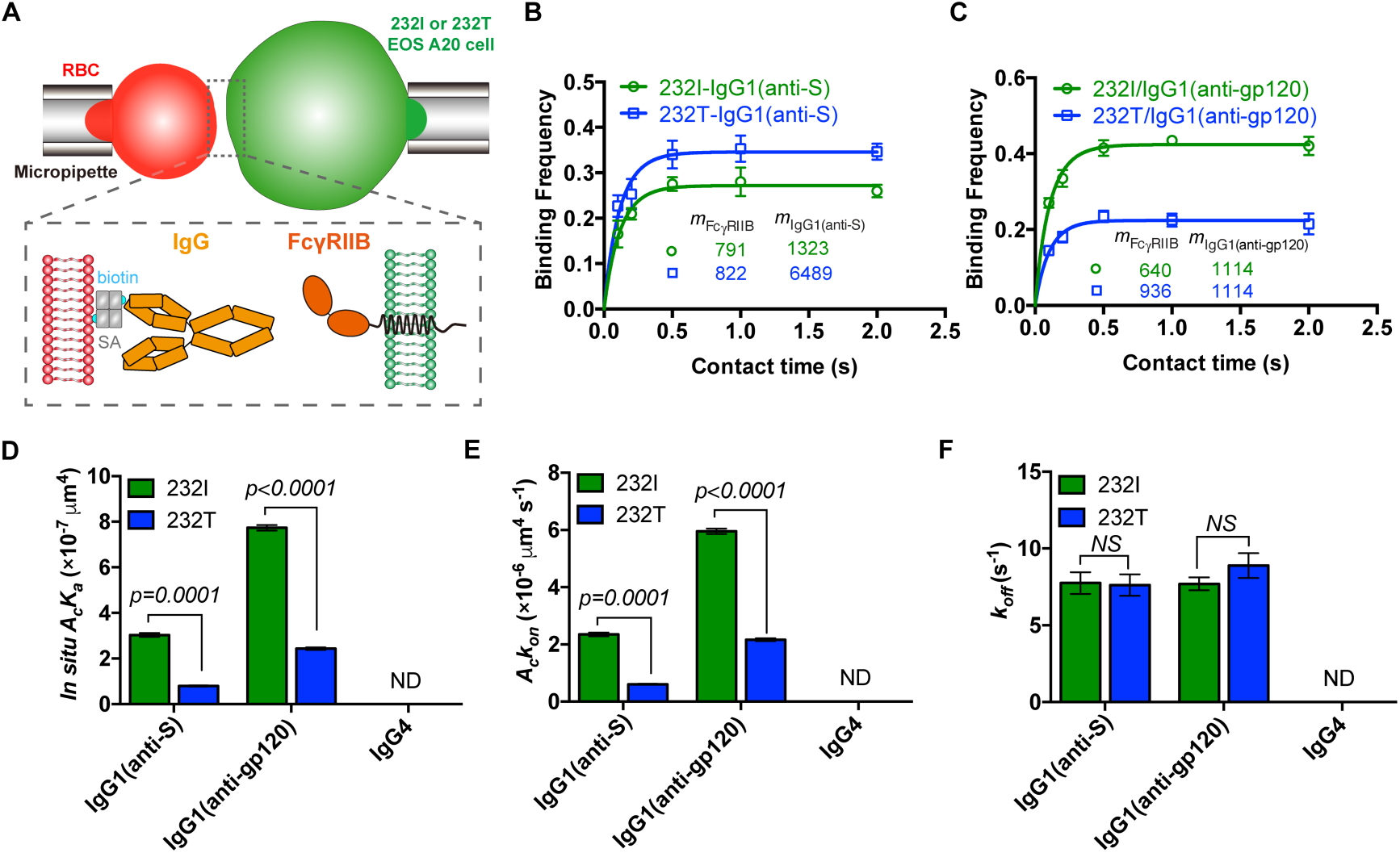
FcγRIIB-232T exhibited significantly reduced 2D IgG1 binding affinity and on-rate in comparison with FcγRIIB-232I. (A) Schematic illustration of micropipette binding frequency approach. Two opposing micropipettes aspirated red blood cell (red, coating IgG antibody) and FcγRIIB A20II1.6 B cell (232I or 232T, green) respectively to operate contact-retraction cycles manipulation. (B and C) Adhesion curves of FcγRIIB (232I or 232T) and human IgG1 antibody (anti-S or anti-gp120) according to probabilistic kinetic model. (D, E and F) From the adhesion curves, *in situ* 2D effective binding affinity (*A*_c_*K*_a_), on-rate (*A*_c_*k*_on_) and off-rate (*k*_off_) were calculated. Error bars represent SEM.

In summary, we confirmed that homozygous FcγRIIB-I232T confers dramatically increased risk of developing more severe clinical manifestations in patients with SLE. The pathological relevant of I232T is caused by the inclination of TM domain which leads to differed conformation of FcγRIIB ectodomain. Ectodomain harboring I232T polymorphism bends towards the membrane such that the Fc binding ability is significantly reduced. The hampered Fc recognition ability of FcγRIIB-I232T results in deficiency on its inhibitory function and thus hyper-activated immune cells, potentially leading to SLE and other immune diseases.

## Methods

### SNP rs1050501 genotyping and statistical analysis

The ethics committee of Peking University People’s Hospital approved this study and informed consents were obtained from each patient and healthy volunteer. There were 711 patients fulfilling the 1997 revised classification criteria of the American College of Rheumatology that enrolled in this study. Healthy volunteers were recruited as controls. 4-8ml peripheral blood was acquired from SLE patients and healthy volunteers. Genomic DNA was extracted from peripheral blood samples using the TIANamp Blood DNA Midi Kit (TIANGEN BIOTECH, Beijing) following the manufacturer’s protocol. The TaqMan Genotyping Assays were applied for genotyping of SNP rs1050501 (TaqMan probe C: 5’-VIC-CGCTACAGCA GTCCCAGT-NFQ-3’, TaqMan Probe T: 5’-FAM-CGCTACAGCA ATCCCAGT-NFQ-3’) (Life technology). Amplification and genotyping analyses were performed using ABI 7300 Real-Time PCR system. Relative quantification of probes levels was calculated (7500 Sequence Detection System Software Version 1.4, ABI). Few samples were genotyped by using primers (forward: 5’-AAGGGGAGCC CTTCCCTCTGTT-3’, reverse: 5’-CATCACCCAC CATGTCTCAC-3’) binding to the flanking introns of exon 5 as reported ^10-11^. The DNA sequencing was done by BGI (Beijing). The Pearson chi-square tests were performed for the comparison of differences between cases and controls at genotype model (recessive model CC vs. TT+TC). The odds ratios (OR), 95% confidence intervals (CI) and p value for recessive model analysis were calculated using logistic regression, adjusting for age and sex. In statistical analyses, *p* value of less than 0.05 was considered statistically significant.

### Molecular Dynamics Simulations

Structure model of the full human FcγRIIB system (residues A46-I310) was built by fusing the crystal structure of the ectodomain (PDB code 2FCB, residues A46-Q215) to the transmembrane (TM) helix (residues M222-R248) model obtained in previous study^16^, the stalk (residues A216-P221) and cytoplasmic regions (residues K249-I310) are randomly placed. An asymmetric lipid bilayer with the membrane lateral area of 100×100 Å^2^ was generated with Membrane Builder in CHARMM-GUI^19^, the lipids in the outer leaflet contain POPC, PSM, and Cholesterol with molar ratio 1:1:1 and these in the inner leaflet contain POPE, POPC, POPS, POPIP2, Cholesterol with molar ratio 4:3:2:1:5. FcγRIIB model was inserted into the lipid membrane with its TM perpendicular to the bilayer surface and the ectodomain stands straight, as shown in Figure 1A.

The WT system was subsequently solvated in 100×100×203 Å^3^ rectagular water boxes with TIP3P water model and was neutralized by 0.15 M NaCl. The I232T polymorphism was obtained from the same configuration using the Mutator plugin of VMD^20^. The final systems contained ∼0.20 million atoms in total.

Both systems were first pre-equilibrated with the following three steps: (1) 5,000 steps energy minimization with the heavy atoms of protein and the head group of the lipids fixed, followed by 2 ns equilibration simulation under 1 fs timestep with these atoms constrained by 5 kcal/mol/Å^2^ spring; (2) 5,000 steps energy minimization with the heavy atoms of protein fixed, followed by 2ns equilibration simulation under 1 fs timestep with these atoms constrained by 1 kcal/mol/ Å^2^ spring; (3) 4 ns equilibration simulation under 2 fs timestep with the heavy atoms of protein ecto- and TM domains constrained (that is, the stalk and intracellular portion is free) by 0.2 kcal/mol/Å^2^ spring.

The resulted systems were subjected to productive simulations for 200 ns with 2 fs timestep without any constrains, and the snapshots of the last 80 ns (sampled at 10 ps intervals) were used for detailed analyses including the probability distributions of hydrogen bonds, tilting angles of the TM helix, inclination angles of ectodomain, the distance between Ig-like C2-type 1 domain and lipid bilayer. The tilting angle of TM helix is defined as the angle between TM helix and membrane plane, similar as that used in previous study^16^. The inclination angle of ectodomain is defined as the angle between the membrane plane and the vector linking NT-terminal of TM helix (M222-I224) and linker region of Ig-like C2-type 1 and 2 domain (S130-W132). The distance between Ig-like C2-type 1 domain and lipid bilayer is defined as the length between center of mass (COM) of this domain and the heavy atoms of phospholipid head in the normal direction of bilayer.

All simulations were performed with NAMD2 software^21^ using CHARMM36m force field with the CMAP correction^22^. The simulations were performed in NPT ensemble (1 atm, 310K) using a Langevin thermostat and Nosé-Hoover Langevin piston method ^23^, respectively. 12Å cutoff with 10 to 12 Å smooth switching was used for the calculation of the van der Waals interactions. The electrostatic interactions were computed using the particle mesh Eward method under periodic boundary conditions. The system preparations and illustrations were conducted using VMD.

### Plasmid construction and cell lines establishment

FcγRIIB-232I pHAGE and FcγRIIB-232T pHAGE plasmids were previously constructed ^16^. mTFP was fused to the N termini of FcγRIIB (232I or 232T) in a pHAGE backbone by ClonExpress(tm) MultiS One Step Cloning Kit (Catalog#C113, Vazyme, China). Stable mTFP-232I/mTFP-232T expressing A20II1.6 B cell lines were acquired by lentivirus infection (three-vector system: mTFP-232I or mTFP-232I pHAGE, psPAX2, and pMD2.G). A20II1.6 B cell lines expressing similar level of either mTFP-232I or mTFP-232T FcγRIIB was obtained by multiple rounds of cell sorting. FcγRIIB-232I and FcγRIIB-232T A20II1.6 B cell lines were previously established^16^.

### FRET measurement

FRET measurements were performed as previously described^24-25^. Briefly, all FRET measurements were carried out on Nikon TiE C2 confocal microscope with 100x oil lens, Argon 457 nm and HeNe 561 nm laser line laser. 1×10^6^ mTFP-232I/mTFP-232T A20II1.6 B cells were stained with 300 nM octadecyl rhodamine B (R18) on ice for 3 mins and then were captured in both channels before or after R18 photo-bleaching. mTFP intensity was processed through Image J. And FRET efficiency= (DQ-Q)/DQ, where DQ and Q are dequenched and quenched mTFP intensity, respectively. FRET efficiency of mTFP-232I or mTFP-232T cells (∼20 cells, respectively) were calculated and plotted through Prism 7. Error bars represent SEM.

### RBC preparation

Streptavidin-coated red blood cells (RBCs) preparation have been described previously ^18^. IgG was biotinylated by EZ-Link Sulfo-NHS-LC-Biotin kits (Thermo Fisher Scientific). Different amounts of biotinylated IgG was linked into RBCs through SA-biotin interaction at RT for 30 min, respectively. IgG-coated RBCs were obtained for micropipette adhesion frequency assay to measure 2D binding kinetics of FcγRIIB/IgG. All above experimental processes were followed by the institutional ethical review board of Zhejiang University.

### 2D binding kinetics measurements

The micropipette adhesion frequency assay was applied to measure FcγRIIB/IgG 2D *in-situ* binding kinetics. The detail experimental progress was previously described^18^. In brief, biotinylated human antibodies (IgG1 or IgG4) were coated on red blood cell (RBC) with streptavidin-biotin association. Two opposing micropipettes aspirated the RBC and FcγRIIB A20II1.6 B cell (232I or 232T) respectively to operate contact-retraction cycles manipulation. Through these 50 contact-retraction cycles, the binding frequency was acquired with definite contact area and a series of setting contact time (0.1, 0.2, 0.5, 1 and 2 s). 3∼4 cell pairs were tested for each setting contact time. And these data were fitted by probabilistic kinetic model. In order to accurately calculate 2D binding affinity and on-rate, these two surface molecular densities (mFcγRIIB and mIgG) were determined by standard calibration beads on flow cytometry, respectively. Binding kinetics were calculated and plotted through Prism 7. Error bars represent SEM.

## Author contributions

W. Chen, J. Lou, and W. Liu conceived this project; W. Chen, J. Lou, W. Liu, W. Hu, Y. Zhang and X. Sun designed the project; W. Hu and W. Chen performed FRET and binding assay; Y. Zhang and J. Lou performed MD simulations; X. Sun and Z. Li performed SNP rs1050501 genotyping and statistical analysis. L. Xu and H. Xie prepared reagents and performed antibody biotinylation. W. Chen, J. Lou, W. Liu, W. Hu, Y. Zhang and X. Sun wrote the manuscript.

## Acknowledgments

We thank Dr. Y. Shi from The Institute of Microbiology of the Chinese Academy of Sciences (IMCAS) for kindly providing us HIV1 gp120 human IgG1 and PD1 human IgG4 antibody, Dr. L. Zhang and X. Wang from Tsinghua University for kindly providing us MERS virus S protein human IgG1 antibody, T. Zhang in W. Chen’s Lab for FRET assay assistance, core facilities in Zhejiang University School of Medicine for technical supports, especially X. Song for FACS supports. This work was supported by grants from the National Basic Research Program of China (2015CB910800 to W. Chen), the National Science Foundation of China (31470900 and 31522021 to W. Chen; 11672317 to J. Lou; 11772348 to Y. Zhang), the Young Thousand Talents Plan of China (W. Chen), the Fundamental Research Funds for the Central Universities (2015XZZX004-32 to W. Chen). The computational resources were provided by the National Supercomputing Center Tianjin Center and HPC-Service Station at the Center for Biological Imaging of the Institute of Biophysics.

---

**Figure 1—figure supplement 1.**
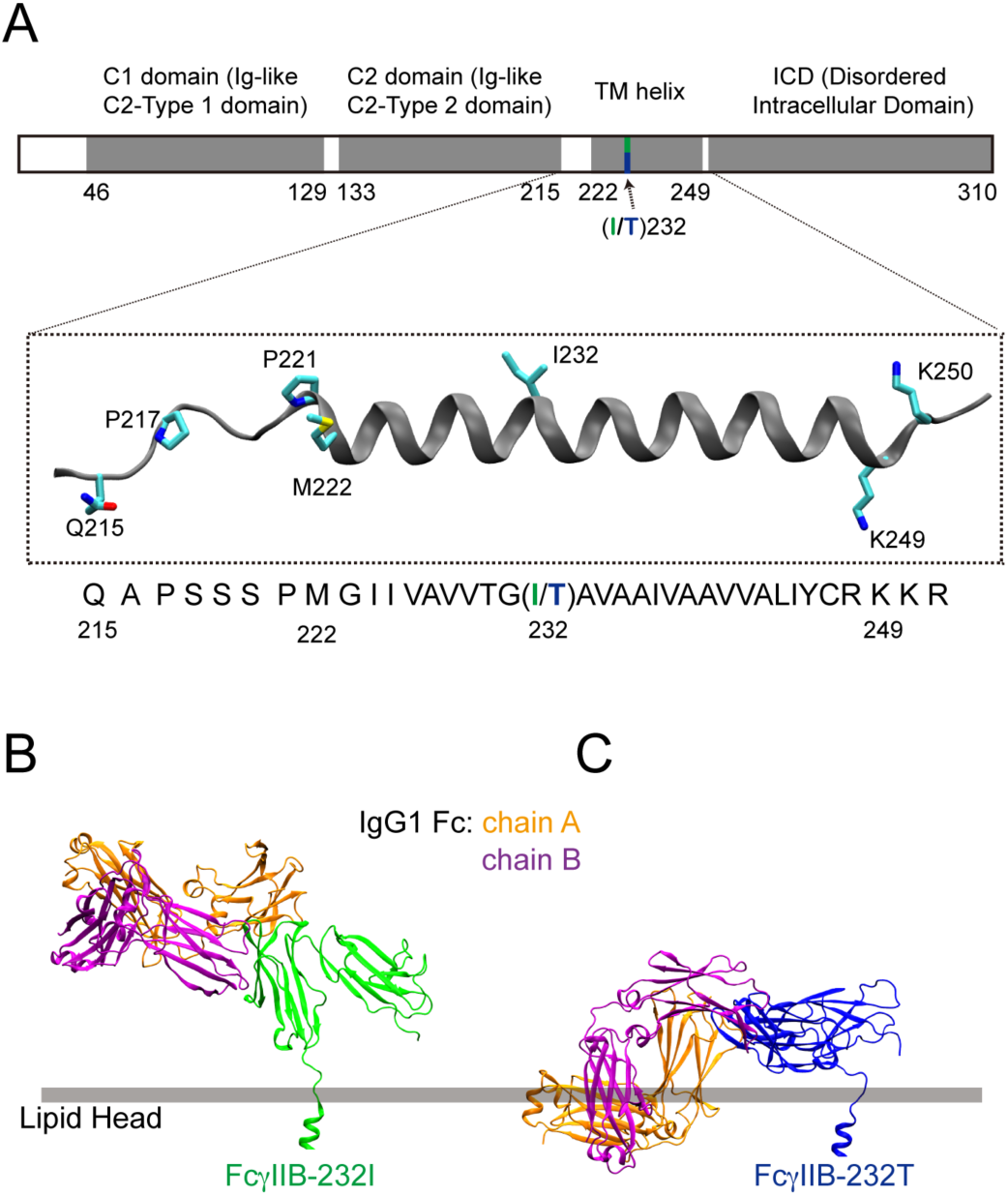
I232T polymorphic change of FcγRIIB induced the recumbent of its ectodomain and may result in impaired binding to Fc domain of antibodies. (A) The domain architecture of FcγRIIB, the modeled structure of the transmembrane region is highlighted, residue 232 is the only difference between WT FcγRIIB (I232) and I232T polymorphism(T232). (B) The ectodomain of WT (FcγRIIB-232I) stands straight with the membrane and is free for antibody biding. (C) For FcγRIIB-232T polymorphic change, although the Fc binding site is still accessible, but its binding ability with antibody will be significantly reduced as observed by the clashes of the Fc domain and the membrane when the ectodomain in the FcγRIIB/Fc complex structure (PDB code: 3WJJ) is superimposed to that observed by the MD simulation.

**Figure 1—figure supplement 2.**
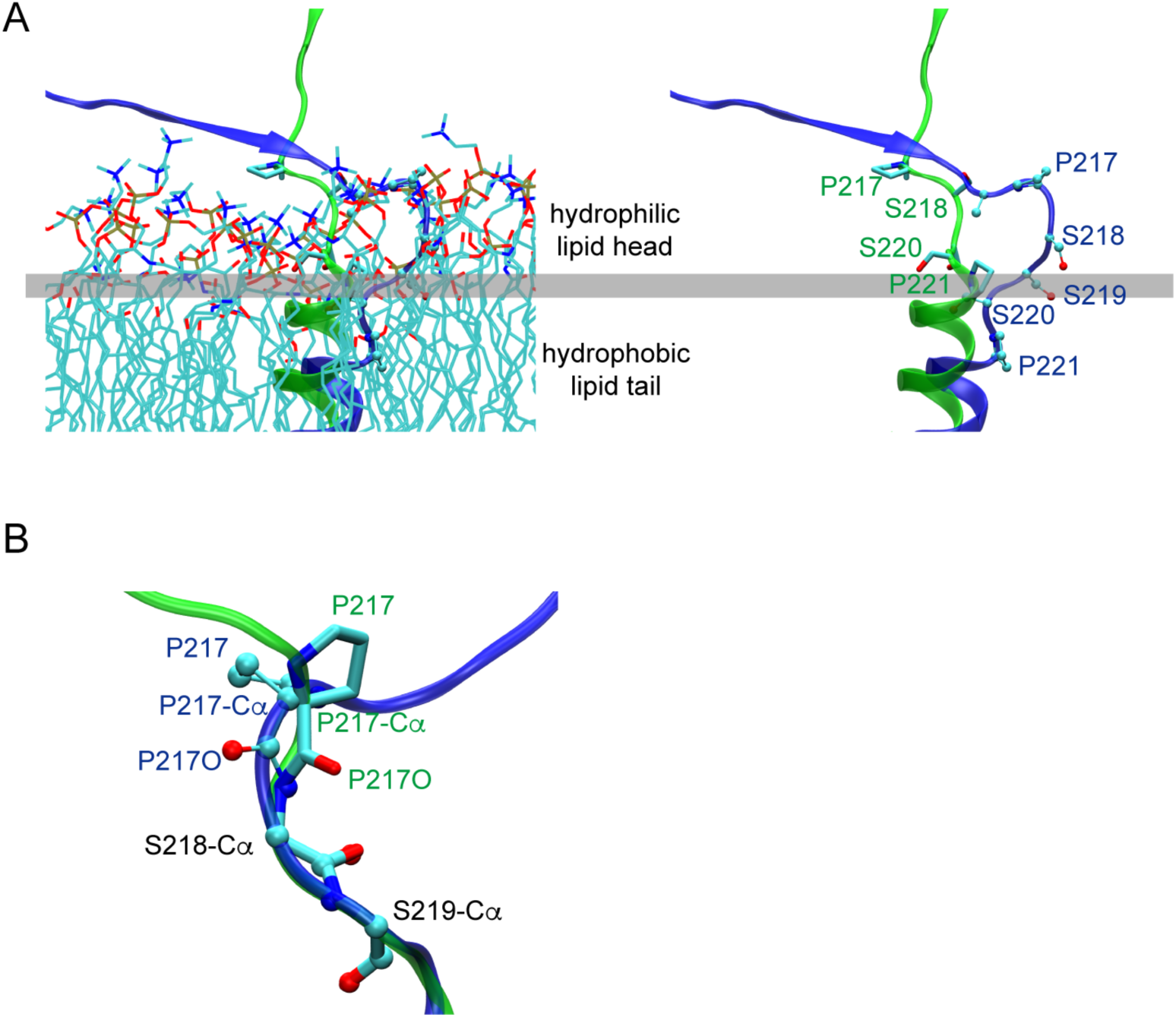
Conformation difference of ecto-TM linker and its vicinity in WT (I232) FcγRIIB and I232T polymorphism obtained by MD simulations. (A)The conformation of the ecto-TM linker in lipid bilayer (left) or lipids removed (right, the residues number in this region is indicated). (B) Conformational difference of I212-S220 regions after aligning residues S218 to S220.

**Figure 1—figure supplement 3.**
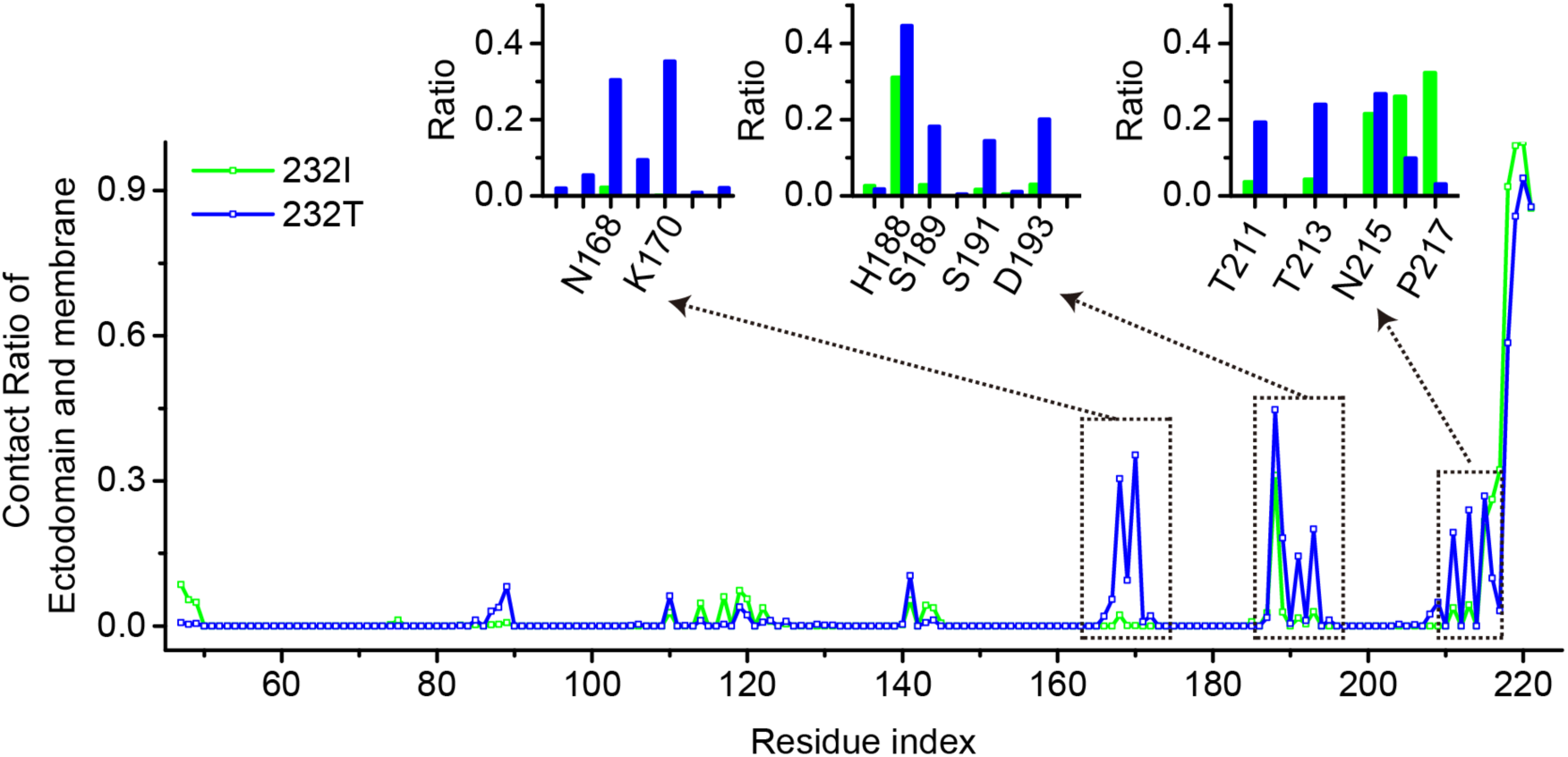
The association between FcγRIIB ectodomain and membrane in I232T polymorphism is mediated by multiple residues. The residues important to the association can be obtained by comparing the contact ratios per residue for WT (232I, green) or the FcγRIIB-232T (232T, blue). Regions with greater contact ration differences are highlighted in the insets. Of them, N168, K170, S191 and D193 may play more essential roles.

